# Arabidopsis ERF109 regulates auxin transport-related genes in root development

**DOI:** 10.1101/725572

**Authors:** Rui Liu, Xiao-Teng Cai, Ping-Xia Zhao, Ping Xu, Cheng-Bin Xiang

## Abstract

The transcription factor ERF109 acts as a crosstalk node between jasmonic acid signaling and auxin biosynthesis by directly regulating *YUC2* and *ASA1* during lateral root formation in Arabidopsis. However, whether ERF109 regulates the auxin transport remains unclear. Here we report a mechanism of ERF109-mediated auxin transport in root system. Through root transcriptome comparison between *erf109*, wild type, and *35S:ERF109*, we found that the genes *PIN2* and *PIN4*, encoding the major membrane-based efflux carriers of auxin, were enriched in the overexpression line. In the promoters of these auxin transport genes, GCC-box cis elements were found and potentially bound by ERF109. Moreover, *PID*, encoding a key regulator in polar auxin transport, was found upregulated in *35S:ERF109* and down regulated in *erf109*. Yeast-one-hybrid and chromatin immunoprecipitation assays showed that ERF109 directly bound to the GCC-box of *PIN2*, *PIN4*, and *PID*. Genetic analyses with double mutants confirmed the function of ERF109 in the regulation of auxin transport in Arabidopsis roots. Taken together, our results show that ERF109 modulates auxin transport by directly regulating *PIN2*, *PIN4* and *PID*. This ERF109-mediated auxin transport likely works together with ERF109-mediated auxin synthesis to establish auxin maxima for lateral root initiation.

## INTRODUCTION

Plant root system is essential for nutrition absorption and mechanical functions. The elaboration of root architecture is crucial for its adaptation to environmental constraints (Ge *et al.*, 2019; Gutierrez-Alanis *et al.*, 2018; Julkowska *et al.*, 2017; Lavenus *et al.*, 2016; Nibau *et al.*, 2008; Scheres and Laskowski, 2016). The phytohormone auxin is well known as a key regulator in plant root configuration and development (Friml, 2003; Matthes *et al.*, 2019; Moller *et al.*, 2017; Moore *et al.*, 2017; Overvoorde *et al.*, 2010; Stoeckle *et al.*, 2018). The Trp-dependent primary auxin biosynthesis and metabolic pathway in Arabidopsis is well established (Zhao, 2012). Accumulating evidence supports that local auxin biosynthesis and its regulation play essential roles in plant development and responses to environmental cues (Zhao, 2018b). The synthesis, transport, and cascade signaling pathways of auxin orchestrate almost every aspect of root development, including primary root (PR) growth, lateral root (LR) initiation, and root tropic growth (Friml *et al.*, 2002b; Fukaki *et al.*, 2007; Miao *et al.*, 2018; Ruzicka and Hejatko, 2017; Spalding, 2013; Weiste *et al.*, 2017; Zhao, 2018a).

Auxin gradient created by polar auxin transport (PAT) in root system is fundamental for plant organogenesis and development (Band *et al.*, 2014; Benkova *et al.*, 2003; Paponov *et al.*, 2005). The active directional cell-to-cell movement of auxin, which is mediated by membrane-based influx and efflux carriers, controls auxin distribution in root system (Brumos *et al.*, 2018; Grigolon *et al.*, 2015; Overvoorde *et al.*, 2010; Petrasek and Friml, 2009). Collectively, there are three major auxin carriers that have been identified in Arabidopsis: PIN-FORMED (PIN) family (auxin efflux carriers), AUXIN RESISTANT 1/LIKE AUX1 (AUX1/LAX) family (auxin influx carriers), ATP-BINDING CASSETTE B (ABCB) family (Carrier *et al.*, 2008; Paponov *et al.*, 2005; Rutschow *et al.*, 2014; Tromas and Perrot-Rechenmann, 2010; Zhou and Luo, 2018).

The PIN family proteins facilitate auxin efflux from the cytoplasm to the apoplast (Adamowski and Friml, 2015; Blilou *et al.*, 2005; Friml, 2010; Keicher *et al.*, 2017; Prat *et al.*, 2018; Ren and Lin, 2015). PIN family has a total of 8 members. Each member has particular function and tissue-specific location. PIN proteins can be classified into the PM-localized group including PIN1–PIN4 and PIN7 that contributing to cell-to-cell auxin transport, and the ER-localized group consisting of PIN5, PIN6 and PIN8 that largely contribute to the regulation of the cellular auxin homeostasis (Wabnik *et al.*, 2011). PIN1 primarily regulates plant meristem tissue and organ development process at the rootward faces of the stele (Huang *et al.*, 2010; O’Connor *et al.*, 2017; Paponov *et al.*, 2005; Xi *et al.*, 2016). PIN2 is observed on the shootward faces of epidermal cells, and regulates root gravitropism through the redistribution of auxin among root tissue (Mendez-Bravo *et al.*, 2019; Muller *et al.*, 1998; Vieten *et al.*, 2005; Zhu *et al.*, 2017). PIN3 concentrates on the basal side of vascular cells as well as the lateral side of pericycle cells in the elongation zone, whose main function is to modulate differential growth and regulate tropic growth (Chen *et al.*, 2015; Friml *et al.*, 2002b; Petrasek and Friml, 2009; Zhang *et al.*, 2013). *PIN4* is observed on outer layer cells and peripheral cells around the quiescent center (QC), and meanwhile has a polar localization in the basal membranes of provascular cells, which regulates both auxin homeostasis and patterning through sink-mediated auxin distribution in root tips (Blilou *et al.*, 2005; Friml *et al.*, 2002a; Xi *et al.*, 2018). PIN7 is detected in the meristem and elongation zone, and located at lateral and basal membranes of provascular cells, which mainly mediates auxin efflux from the QC area and regulates root tropism and early embryonic development (Band *et al.*, 2014; Blilou *et al.*, 2005; Ki *et al.*, 2016; Ruiz Rosquete *et al.*, 2018). The ER-localized PIN5, PIN6 and PIN8 do not transport auxin from cell to cell but regulates intracellular auxin homeostasis and development (Cazzonelli *et al.*, 2013; Ding *et al.*, 2012; Mravec *et al.*, 2009).

AUX1/LAX membrane proteins were shown to play a significant role in the regulation of PAT. There are four members (AUX1, LAX1, LAX2 and LAX3) in Arabidopsis genome, which facilitate auxin influx from the apoplast to the cytoplasm (Robert *et al.*, 2015; Rutschow *et al.*, 2014; Swarup and Peret, 2012). AUX1 mainly regulates root hair development and gravitropism (Dindas *et al.*, 2018; Street *et al.*, 2016; Wakeel *et al.*, 2018). Many researches have implicated that AUX1 and LAX3 can regulate lateral root development and apical hook formation (Jones *et al.*, 2009; Porco *et al.*, 2016; Swarup *et al.*, 2005; Swarup and Peret, 2012; Wakeel *et al.*, 2018).

Members of the ABCB family are also involved in auxin transportation (Cho and Cho, 2013; Kaneda *et al.*, 2011; Zhang *et al.*, 2018). Some other classes of membrane protein, such as potassium and nitrate transporters, have been reported to play a significant role in facilitating auxin movement as well (Krouk *et al.*, 2010; Vicenteagullo *et al.*, 2004).

The PIN family proteins can be specifically regulated by a group of kinases, which includes serine/threonine kinase PINOID (*PID*), PID2 and WAVY ROOT GROWTH (WAG) proteins (Barbosa *et al.*, 2018; Grones *et al.*, 2018; Haga *et al.*, 2014; Weller *et al.*, 2017). Loss-of-function mutants of the *PID* display apical organogenesis defects and the decline of auxin transport, which is similar to the pin1 mutant (Benjamins *et al.*, 2001; Bennett *et al.*, 1995). The overexpression of *PID* causes the reduction of lateral auxin transport due to the preferential stimulation of downward directed PAT to the root apex by *PID*, which leads to the loss of auxin of root apex and peripheral cells, and subsequently results in the reduction of lateral roots, shortened hypocotyl and root, and the loss of gravitropism and phototropism (Benjamins *et al.*, 2001; Haga and Sakai, 2015; Saini *et al.*, 2017a; Sukumar *et al.*, 2009; Willige and Chory, 2015). Recent studies have revealed that *PID* is a crucial contributor of the regulation of apical-basal PIN polarity (Friml *et al.*, 2004; Kleine-Vehn *et al.*, 2009; Saini *et al.*, 2017b; Wang *et al.*, 2019; Zourelidou *et al.*, 2014). The mechanisms of *PID*-mediated PIN polarity alteration have been unraveled (Christensen *et al.*, 2000; Dory *et al.*, 2018; Geldner *et al.*, 2003; Kleine-Vehn *et al.*, 2009; Michniewicz *et al.*, 2007).

We previously reported a drought resistance mutant edt1 that displayed an improved root system with more lateral roots, which was caused by an activated expression of the homeodomain-START transcription factor EDT1/HDG11 (Yu *et al.*, 2008). We later found that the phytohormone jasmonic acid (JA) was increased in edt1 roots, consistent with root transcriptome data that show a number of JA biosynthesis-related genes are directly upregulated by EDT1 (Cai *et al.*, 2015b). Moreover, we identified the JA-responsive transcription factor ETHYLENE RESPONSE FACTOR 109 (ERF109) from the same transcriptome analysis. ERF109 is a plant specific transcription factor from the ERF family and acts as an important crosstalk node integrating JA signaling into auxin biosynthesis (Cai *et al.*, 2014). ERF109 is responsive to JA signaling and promotes auxin biosynthesis by binding to the promoters of *YUC2* and *ASA1*, and therefore affects lateral root formation. As a typical ERF family protein, ERF109 contains one APETALA2 (AP2) domain and can directly bind to the GCC-box (5’-GCCGCC-3’) in the promoters of its target genes to regulate their expressions (Barczak *et al.*, 2016; Cai *et al.*, 2014; Fujimoto *et al.*, 2000; Matsuo *et al.*, 2015). Meanwhile, ERF109 is responsive to a variety of biotic and abiotic redox stimulation and thus also named as Redox-responsive Transcription Factor 1 (RRTF1) (Khandelwal *et al.*, 2008; Matsuo *et al.*, 2015; Matsuo and Oelmüller, 2015; Wang *et al.*, 2008). Recent reports show that ERF109 is also involved in the activation of root stem cells under JA signal (Zhou *et al.*, 2019) and plant regeneration promoted by jasmonate-mediated wound signaling (Zhang *et al.*, 2019).

We previously noticed the reduction of auxin level in root tip of loss-of-function mutant *erf109*. However, ERF109 is not expressed in primary root tip under normal conditions (Cai *et al.*, 2014), which strongly implicates that ERF109 may also regulate auxin transport. Here we report that ERF109 can also up regulate the transcript level of auxin transport-related *PIN2*, *PIN4*, and *PID*. Yeast-one-hybrid and ChIP-qPCR assays demonstrated that ERF109 can directly bind to GCC-boxes of *PID*, *PIN2* and *PIN4* to up regulate their expression, which is supported by genetic analyses. Our results demonstrate that ERF109 modulates lateral root formation by regulating auxin transport in addition to auxin biosynthesis we previously reported.

## RESULTS

### ERF109 affects the expression of auxin transport genes

To investigate whether ERF109 regulates auxin transport, we first confirmed the expression of ERF109 and the phenotype of the *erf109* mutant, *ERF109* overexpression line with the wild type (Figure 1A and B). Then we measured the transcript level of auxin transport-related genes in the root tissue by quantitative RT-PCR. The results showed that the transcript level of *PIN1* was decreased in *erf109* mutant, but not increased in *ERF109* overexpression line significantly (Figure 1C). The transcription of *PIN2* and *PIN4* was significantly increased in *35S:ERF109*, but did not show obvious reduction in *erf109* mutant (Figure 1D and F). There is no significant change in the transcript levels of *PIN3*, *PIN7* and *AUX1* in *erf109* and *35S:ERF109* compared with that in wild type (Figure 1E, G and H).

**Figure 1.**
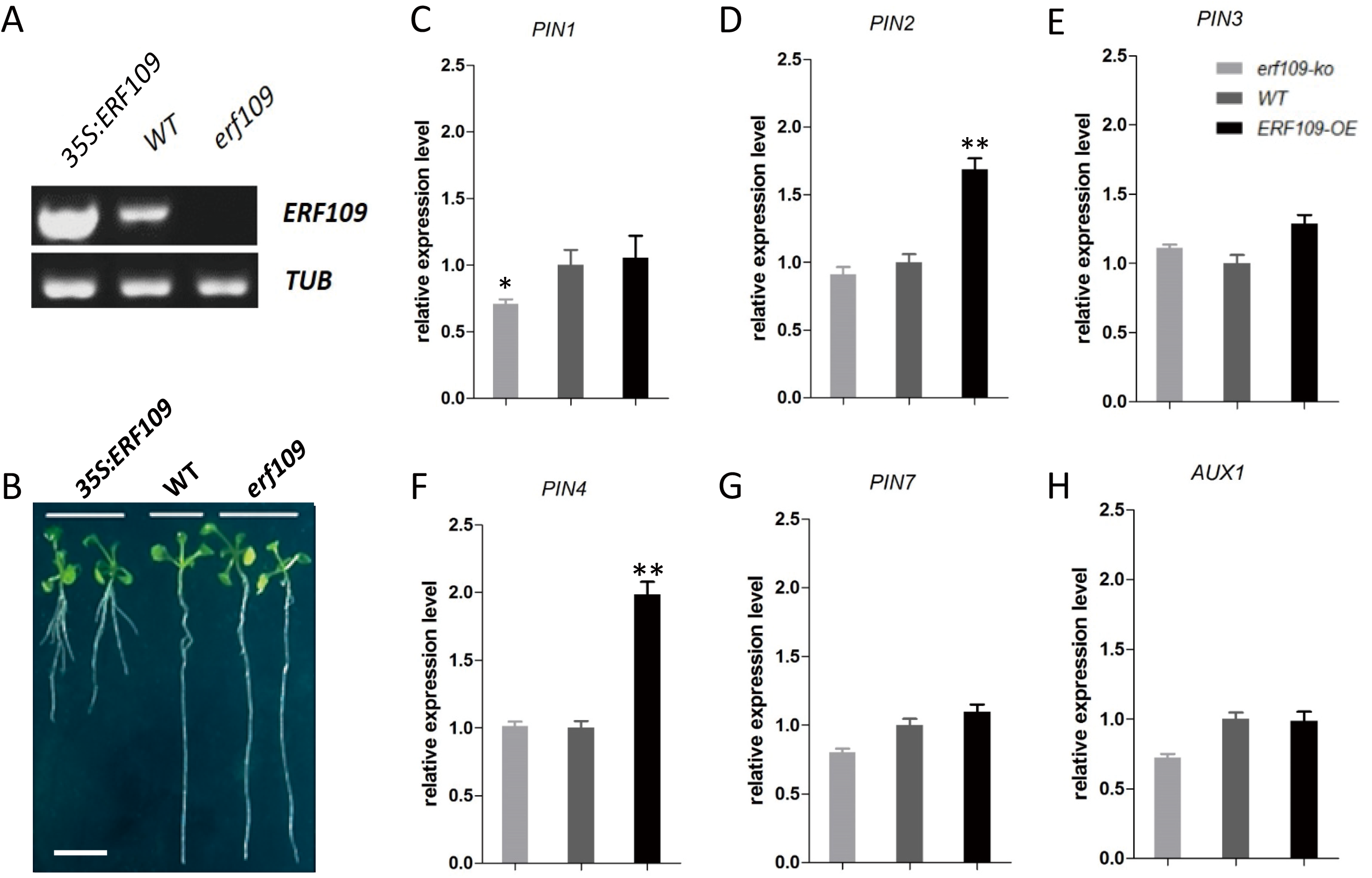
ERF109 regulates the expression of auxin transport genes. **A**. Confirming the expression level of *ERF109* by RT-PCR. RNA samples isolated from 10-d-old seedlings of *erf109*, *35S:ERF109* and wild type plants were analyzed by quantitative RT-PCR. **B**. Confirming the phenotype of 7-day-old *35S:ERF109* plants. *35S:ERF109* seedlings had shorter primary root and more lateral roots than the wild type. Scale bar = 1 cm. **C-H**. Quantitative RT–PCR analysis of the transcript levels of auxin transport genes. Transcript level of *PIN1 (C)*, *PIN2 (D)*, *PIN3 (E)*, *PIN4 (F)*, *PIN7 (G)*, and *AUX1 (H)* in the wild type, *erf109*, and *35S:ERF109* seedlings. RNA was isolated from the two-week-old seedlings. Values are mean ± SD (n=3 experiments, *P<0.05, **P<0.01). Asterisks indicate Student’s t-test significant differences.

We also observed the protein level and localization pattern of auxin transporters by using GFP reporter lines in *35S:ERF109* and wild type background. Both *PIN2*:GFP and *PIN4*:GFP reporter showed higher GFP signals in the *35S:ERF109* line compared with that in the wild type, but the localization pattern of *PIN2* and *PIN4* was unchanged (Figure 2). Other auxin transport proteins had no significant change in the expression level or localization pattern as shown by the reporter lines *PIN1pro:PIN1:GFP, PIN3pro:PIN3:GFP, PIN7pro: PIN7:GFP, AUX1pro:AUX1:GFP* in the wild type and *35S:ERF109* background (Supplemental Figure S1).

**Figure 2.**
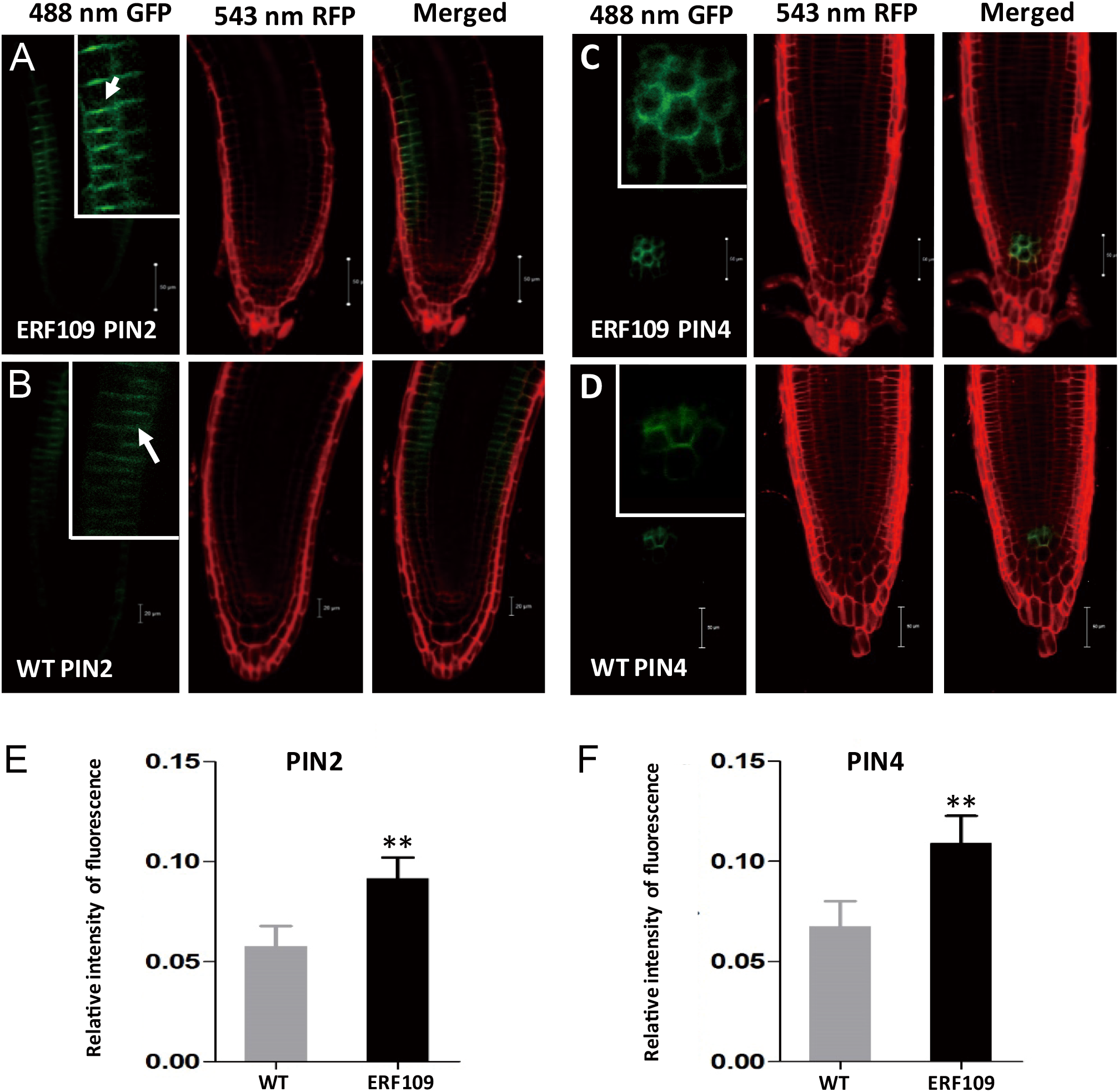
Fluorescence intensity of PIN2-GFP and *PIN4*-GFP was elevated in ERF109 over-expression line. PIN2-and PIN4-GFP reporters were introduced in the wild type and *35S:ERF109* by genetic crosses [*35S:ERF109f* ♂ *× PIN2pro:PIN2:GFP*♀ (ERF109 PIN2, A), wild type ♂ ×*PIN2pro:PIN2:GFP*♀ (WT PIN2, B), *35S:ERF109f* ♂ *× PIN4pro:PIN4:GFP*♀ (ERF109 PIN4, C), and wild typef ♂ × *PIN4pro:PIN4:GFP*♀ (WT PIN4, D)]. The GFP signals were observed using 7-day-old seedlings on a confocal microscope. Bar=50 μM. Three independent lines were analyzed for wild type and *35S:ERF109* background, and consistent results were obtained. GFP intensity of PIN2 (E) and PIN4 (F) was quantified by Image J (https://imagej.nih.gov/ij/). Values are mean ± SD (n=3 experiments, **P<0.01). Asterisks indicate Student’s t-test significant differences.

### ERF109 directly binds to the GCC-boxes in the promoter of *PIN2* and *PIN4*

Considering that ERF109 is a transcription factor, we speculate that ERF109 may transcriptionally regulate the expression of *PIN2* and *PIN4*. To identify whether these auxin transport genes are potential targets of ERF109, we searched the GCC-boxes in the promoters and transcribed region of these auxin transport genes and found that *PIN2* and *PIN4* contained GCC-box. *PIN2* has one GCC-box in the promoter and two GCC-boxes in its transcribed region. *PIN4* contains two GCC-boxes in the transcribed region (Figure 3A).

**Figure 3.**
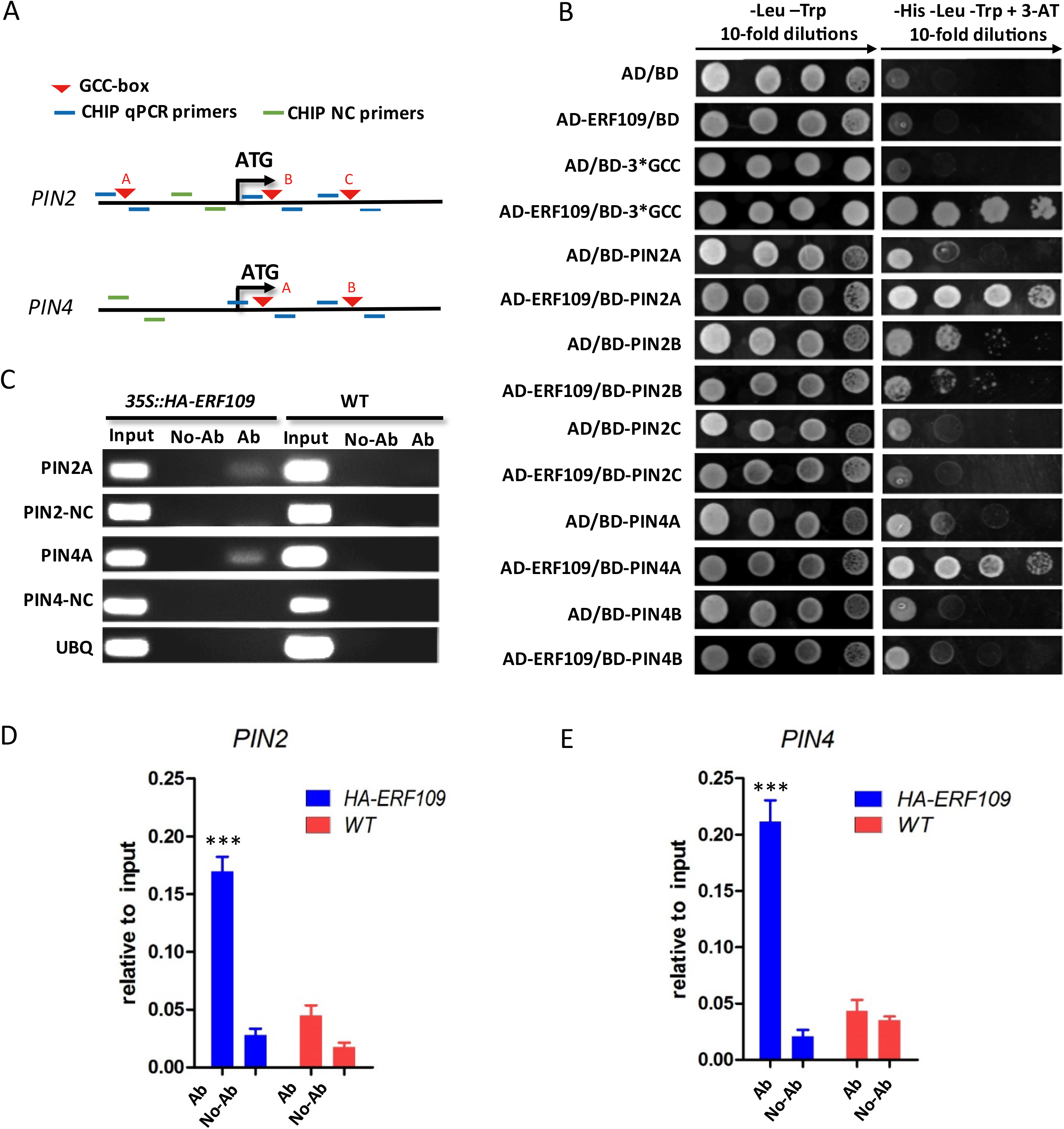
ERF109 directly regulates the transcription of *PIN2* and *PIN4*. **A**. The schematic view of the locations of GCC-boxes (inverted red triangles) in the promoters and transcription regions of *PIN2* and *PIN4* as well as the primers used for ChIP-PCR (blue lines for fragments contained GCC-box and green lines for fragments without GCC-box worked as negative control). **B**. Yeast-one-hybrid assay for ERF109 binding to the GCC-box of *PIN2* and *PIN4*. 30bp fragments contained GCC-box were chosen for BD vector constructions. A serial yeast dilutions (1:1, 1:10, 1:100 and 1:1000) were grown on SD medium lacking Leu and Trp (SD-Leu-Trp) and transferred to SD medium lacking Leu, Trp and His + 3-AT (SD-Leu-Trp-His) respectively. The empty AD (pGADT7) and BD (pHIS2) vectors were used as negative controls. **C**. ChIP-PCR assay for the binding between ERF109 and GCC-boxes of *PIN2* and *PIN4*. DNA fragments with HA-ERF109 were precipitated from input DNA with anti-HA antibodies or with no antibody. The enrichment of DNA fragments was determined by PCR. The regions of UBQ, *PIN2* and *PIN4* that do not contain GCC-box were used as negative controls. **D-E**. Quantitative PCR analysis for ChIP assay. The enrichments of *PIN2* (D) and *PIN4* (E) fragments contained GCC-box were confirmed by Quantitative-PCR. Values are mean ± SD (n=3 experiments, ***P<0.001). Asterisks indicate Student’s t-test significant differences.

We performed yeast-one-hybrid assay to determine whether ERF109 could directly bind to the GCC-boxes of *PIN2* and *PIN4*. 30bp DNA fragments contained GCC-box in the middle were chosen from both the promoter and transcribed region of *PIN2* and transcribed region of *PIN4*. The fragments A from *PIN2* and *PIN4* displayed high affinities to ERF109 (Figure 3A and B). The results show that ERF109 is able to bind to the GCC-boxes of *PIN2* and *PIN4* in yeast cells.

To confirm whether the interaction between ERF109 and the GCC-boxes of *PIN2* and *PIN4* takes place *in planta*, chromatin immunoprecipitation (ChIP) assays were performed using HA-ERF109 transgenic plants with anti-HA antibodies. These DNA fragments enriched by anti-HA antibodies can be detected by PCR (Figure 3C). Further analysis with quantitative PCR also confirmed the enrichment of GCC-box fragments of *PIN2* and *PIN4* by ERF109 (Figure 3D and E). As the ChIP-PCR and ChIP-qPCR results shown in Figure 3C-E, the *PIN2* and *PIN4* fragments containing the GCC-boxes were significantly enriched by ERF109, which is consistent with yeast-one-hybrid assay results. Thus, the specific binding of ERF109 to the GCC-boxes of *PIN2* and *PIN4* was confirmed in Arabidopsis.

### *PID* is another target of ERF109

Since PIN family auxin transporters can be phosphorylated by a group of kinases, including serine/threonine kinase PINOID (*PID*), PID2 and WAVY ROOT GROWTH (WAG) proteins, we examined the transcript level of these genes and found that the expression of *PID* was down regulated in *erf109* (*erf109-ko*) and up regulated in *35S:ERF109* (ERF109-OE) compared with that in the wild type (Figure 4A). *PID* has a GCC-box in its promoter and two GCC-boxes in transcribed region (Figure 4B). To find out whether these GCC boxes can be bound by ERF109, we conducted yeast-one-hybrid assays and found that ERF109 was indeed able to bind to the *PID* B cis element (Figure 4C). To confirm whether this specific binding occurs in planta, we performed ChIP assays and found that the *PID* B fragment containing the GCC-box was significantly enriched by ERF109 in both ChIP-PCR and ChIP-qPCR (Figure 4D and E).

**Figure 4.**
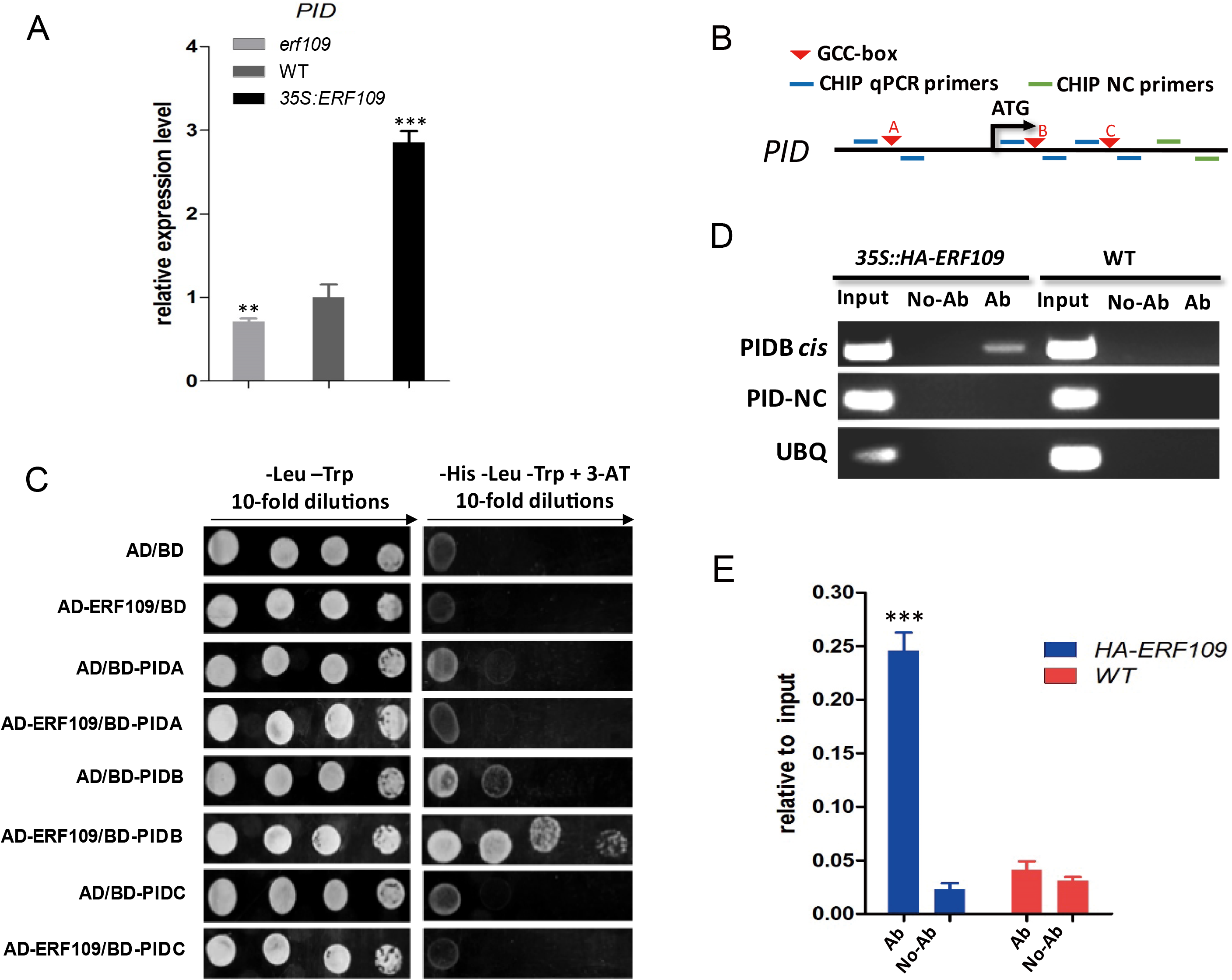
*PID* is another direct target of ERF109. **A**. Expression levels of *PID* by quantitative RT-PCR. RNA samples were isolated from 7-d-old seedlings of *erf109* knockout mutant (*erf109* ko), wild type (WT), and the overexpression line *35S:ERF109* (ERF109-OE). Values are mean ± SD (n=3 experiments, **P<0.01, ***P<0.001). Asterisks indicate Student’s t-test significant differences. **B**. The schematic view of the location of GCC-box cis-element (inverted red triangle) in the promoter and transcribed region of *PID* as well as the primers used for PCR (blue lines for fragments contained GCC-box and green lines for fragments without GCC-box worked as negative control). **C**. Yeast-one-hybrid assay for ERF109 binding to the GCC-box cis-element of *PID*. The assay was performed as same as figure 3B. **D**. ChIP-PCR assay was conducted using 35S:HA-ERF109 transgenic seedlings and anti-HA antibodies. The assay was performed as for Figure 3C. **E**. Quantitative PCR analysis for ChIP assay. The enrichment of *PID* promoter fragment was confirmed by quantitative PCR. Values are mean ± SD (n=3 experiments, ***P<0.001). Asterisks indicate Student’s t-test significant differences.

### Genetic analyses of ERF109 function in *pin2*, pin4 and pid mutant background

To detect whether *PIN2*, *PIN4* and *PID* function as target genes of ERF109, *35S:ERF109 pid*, *35S:ERF109 pin2* and *35S:ERF109* pin4 were generated by crosses (Supplemental Figure S2) and their root phenotypes were observed. Genetic assay results showed that the primary root length and lateral root number of *35S:ERF109 pin2*, *35S:ERF109 pin4* and *35S:ERF109* pid were intermediate between male parent and female parent (Figure 5). These results demonstrate that *PIN2*, *PIN4* and PID are each partially responsible for the root phenotype of *35S:ERF109*, indicating that *PIN2*, *PIN4*, and *PID* are targets of ERF109.

**Figure 5.**
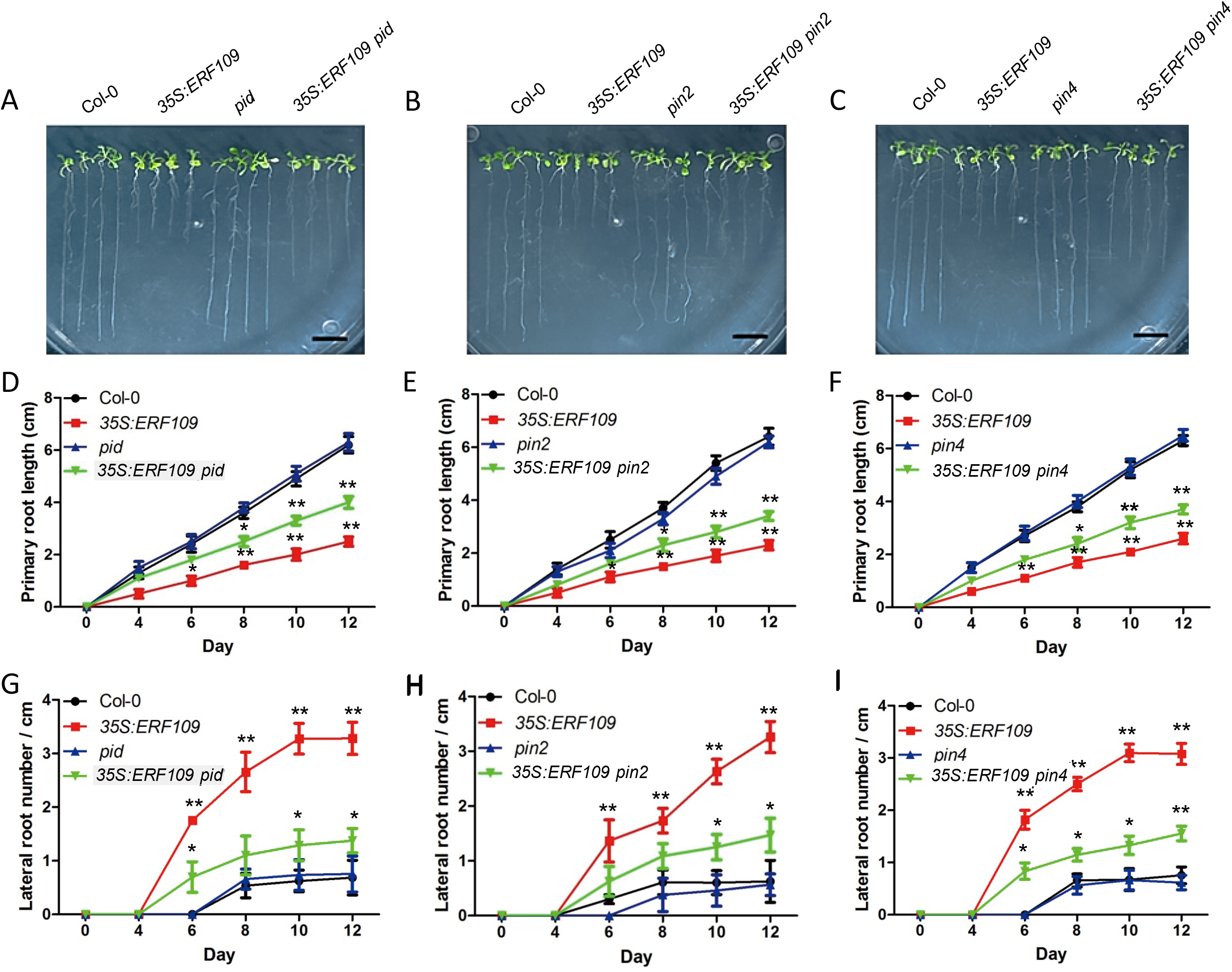
Genetic analyses of ERF109 in pid, *pin2* and pin4 mutant background. **A-C**. Root phenotype. Seeds of indicated lines were separately germinated on MS for 12 days before photographs were taken (Scale bar =1cm). **D-F**. The primary root length was measured at the indicated time points. Values are mean ± SD (n=30 seedlings, *P<0.05, **P<0.01). Statistically significant differences were calculated based on the Student’s t-tests. **G-I**. The numbers of LRs (per cm) of different plants grown on MS were counted at the indicated time points. Values are mean ± SD (n=30 seedlings, *P<0.05, **P<0.01). Statistically significant differences were calculated based on the Student’s t-tests.

## DISCUSSION

Plant root system is characterized by its flexibility and adaptability, which allows the optimal adjustment according to environmental constraints (Moore *et al.*, 2017; Overvoorde *et al.*, 2010; Vieten *et al.*, 2005). Both auxin biosynthesis and transport are involved in the regulation of Arabidopsis root development (Du and Scheres, 2018; Morffy and Strader, 2018; Xu *et al.*, 2017). JA has been long recognized as a stress hormone, and can alter root architecture dramatically, but our knowledge about the underlying mechanisms remains limited (Ku *et al.*, 2018; Wasternack, 2007; Wasternack and Hause, 2013). Previous studies have showed that JA can be induced by a variety of environmental stimuli (Carvalhais *et al.*, 2015; de Ollas *et al.*, 2013; Gundlach *et al.*, 1992; Wasternack, 2007). The transcription factor EDT1/HDG11 up-regulates JA biosynthesis and thus improves Arabidopsis lateral root formation (Cai *et al.*, 2015a; Yu *et al.*, 2008). ERF109 is responsive to JA signaling and integrates JA signal into auxin biosynthesis (Cai *et al.*, 2014). In this study, we further unraveled the molecular mechanisms by which JA-responsive ERF109 regulates auxin transport in addition to auxin biosynthesis by directly binding to the GCC boxes of *PIN2*, *PIN4*, and *PID* and activating their expression. The increased *PID* stimulates PIN activities. Collectively, auxin transport is enhanced.

### ERF109 affects auxin transport by directly regulating *PID*, *PIN2* and *PIN4*

Polar auxin transport is mainly mediated by PIN efflux carriers family and AUX1 influx carriers family (Band *et al.*, 2014; Peret *et al.*, 2012; Petrasek and Friml, 2009; Vieten *et al.*, 2005). The transcript level of *PIN2* and *PIN4* increased about 2-fold in *35S:ERF109* compared with that of wide type (Figure 1D, F). Given that ERF109 can directly bind to the GCC-boxes cis-element, it was found that both *PIN2* and *PIN4* contained GCC-box (Figure 3A). Through the Y1H and ChIP assays, ERF109 can bind to the GCC-box cis-element of *PIN2* and *PIN4 in vitro* and *in vivo* (Figure 3B-E). Further genetic assay also confirmed that *PIN2* and *PIN4* were responsible for the phenotype of *35S:ERF109* (Figure 5). However, there is no significant down-regulated expression in *erf109* mutant (Figure 1D and F). Considering that the *ERF109* and *PIN2*, 4 have different expression pattern and localization (Cai *et al.*, 2014; Paponov *et al.*, 2005; Wisniewska *et al.*, 2006), *PIN2* and *PIN4* can be only altered under ectopic expression of ERF109. Notably, we found a significant decline of the expression of PIN1 in the *erf109* line (Figure 1C). Due to that ERF109 mediate auxin biosynthesis in shoot and root tissues (Cai *et al.*, 2014) and PIN genes can be regulated by auxin itself (Prat *et al.*, 2018; Vieten *et al.*, 2005), it is possible that the down-regulated expression of *PIN1* is caused by the decline of auxin level in *erf109* seedlings.

PID is a positive regulator of auxin efflux (Benjamins R, et al. 2001, Lee SH, et al. 2006, Zourelidou M, et al. 2014). It is notable that the *ERF109* and *PID* have similar expression pattern. Both of them are expressed in neonatal vascular tissue, root tip meristem, root primordial and weakly expressed in plant root tips (Benjamins R, et al. 2001, Lee SH, et al. 2006). *PID* was significantly down-regulated in the *erf109* seedlings and up-regulated in *35S:ERF109* lines (Figure 4A). The GCC-box from the *PID* displayed a high affinity to ERF109 (Figure 4B-E) and subsequent genetic experiments confirmed that *PID* is a downstream target of ERF109 (Figure 5A, D and G). Taken together, ERF109 can regulate *PID* by directly binding to the GCC cis-element in the transcribed region. It is conceivable that an activated expression of PID will enhance the phosphorylation of PIN2 and PIN4, leading to enhanced auxin transport.

### A working model for ERF109-regulated auxin homeostasis in roots

Based on our results, we propose a working model for the JA-responsive ERF109 to regulate auxin homeostasis in roots (Figure 6). In this model, ERF109 responds to JA signaling. As a transcription factor, ERF109 directly binds to the GCC-boxes in the promoters of *ASA1* and *YUC2* to promote auxin biosynthesis as we previously showed (Cai *et al.*, 2014). *ASA1* is a key gene required for auxin biosynthesis under JA signal in lateral root development (Sun *et al.*, 2009). On the other hand, ERF109 can also directly bind to the GCC-box of *PID*, *PIN2* and *PIN4* to increase their transcript levels, thus altering auxin transport. These results suggest that both auxin synthesis and transport may work together to generate the auxin gradient in lateral root formation in response to JA.

## MATERIALS AND METHODS

### Materials and growth conditions

*Arabidopsis thaliana* ecotype Columbia-0 (Col-0) was used as wild type. Homozygous loss-function *erf109* mutant, *35S:ERF109* and *35S:HA-ERF109* lines were previously described (Cai *et al.*, 2014). *PINpro:PIN:GFP* lines and *AUX1pro:AUX1:GFP* were used as female parents in genetic analysis, while Col-0 and *35S:ERF109* were used as male parents. The *pid* (CS9421), *pin2* (CS8058), *pin4-3* mutants were ordered from Arabidopsis Biological Resource Center (ABRC).

**Figure 6.**
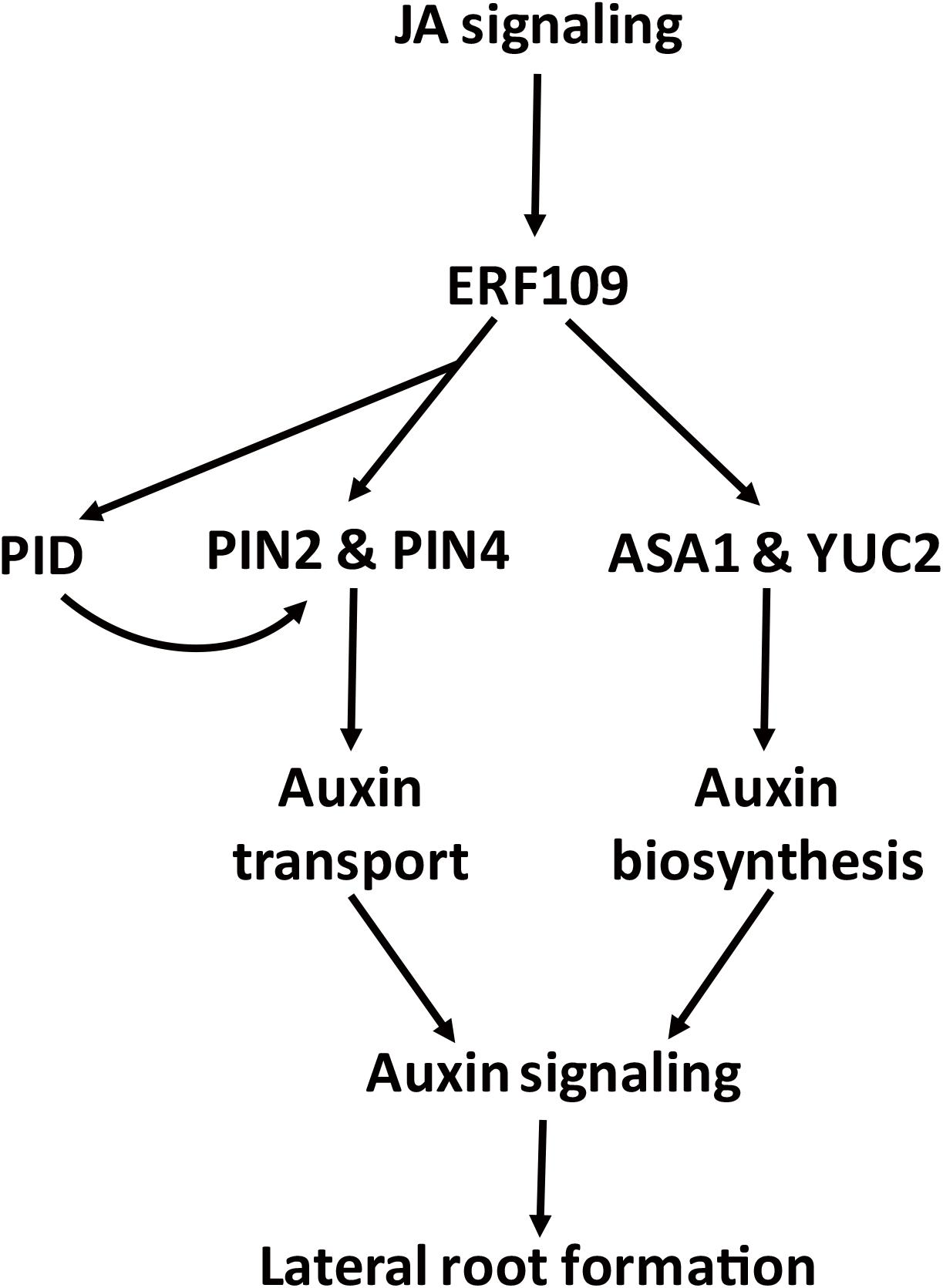
A working model of ERF109 in auxin homeostasis. JA-responsive ERF109 upregulates auxin biosynthesis by directly binding to GCC-boxes in the promoters of *ASA1* and *YUC2*. ERF109 also directly bind to the GCC-boxes cis-elements in *PIN2*, *PIN4*, and *PID*. Elevated *PID* enhances PINs phosphorylation and stimulates auxin transport. Therefore, ERF109 regulates root development through both auxin biosynthesis and transport.

Surface sterilization of Arabidopsis seeds was conducted with 10% bleach for 10 min and seeds were washed five times in sterile environment subsequently. After keeping in darkness for 3 days at 4°C, sterilized seeds were transferred and germinated on vertical 1/2 Murashige and Skoog (MS) medium agar plates (1% (w/v) sucrose) and grown in 22°C under 16-h light/8-h dark cycles.

### Gene expression analysis

Gene expression analysis was conducted by RT-PCR and quantitative RT-PCR. Total RNA was isolated with TRIZOL reagent (Invitrogen) from 10-day-old Col-0, *erf109* and *35S:ERF109* seedlings. Then, cDNA was generated through Reverse transcription reaction with Prime Script RT reagent Kit (TaKaRa). After PCR amplification of cDNA, PCR products were examined on a 2% agarose gel stained with ethidium bromide (EB). Quantitative RT-PCR was carried out with the SYBR Premix Ex Taq II (TaKaRa) on Applied Biosystems Step One Real-Time PCR system. Experiment data was normalized to *Ubiquitin5* (*At3g62250*). The specific primers used in the examination of transcript levels of target genes were listed in Supplemental Table S1. The results of PCR were also examined on a 2% agarose gel stained with EB.

### Confocal microscopy analysis

The GFP observation of root tissues with ZEISS710 confocal laser scanning microscope was described previously (Cai *et al.*, 2014). The GFP fluorescence intensity was quantified by Open Source software Image J (https://imagej.nih.gov/ij/) (Collins, 2007).

### Bioinformatic analysis of GCC-box cis-elements

The GCC-box cis-elements were searched using Arabidopsis cis-regulatory element database (AtcisDB, http://arabidopsis.med.ohio-state.edu/AtcisDB/) (Yilmaz *et al.*, 2011). GCC-box cis-elements were searched in the promoter and transcription region of auxin transport related genes subsequently using TAIR - the Arabidopsis information resource (http://www.arabidopsis.org) for confirmation and supplement of the AtcisDB results.

### Yeast-one-hybrid assay

Yeast-one-hybrid assay (Y1H) was previously described (Cai *et al.*, 2014). The effector plasmid pAD-ERF109 was generated by plasmid pAD-GAL4-2.1 (AD vector) with ERF109-coding region sequence cloned into it, which was used for the expression of fusion protein for DNA binding. The 30bp DNA fragments containing GCC-boxes from *PID*, *PIN2*, and *PIN4* were synthesized (Sangon of Shanghai, China) (Supplemental Table S1) with cohesive ends at each side. Then, these DNA fragments were annealed and ligated into reporter plasmid pHIS2 (BD vector) respectively. The plasmid pHIS2-3*GCC-boxes was also generated by pHIS2 vector with three tandem repeat of GCC-box element. The empty vector of pAD and pHIS2 were used as negative control. Combinations of different AD and BD vectors were co-transformed into Y187 yeast strains. Then, Y187 strains were grown on SD/-Trp-Leu medium and transferred to SD/-Trp-Leu-His medium with 10 mM 3-aminotriazole (3-AT, Sigma) in gradient dilutions (1:10, 1:100 and 1:1,000). The growth state of yeast on SD/-Trp-Leu-His medium indicated the interaction between ERF109 and the corresponding cis-element.

### ChIP assay

Chromatin immunoprecipitation (ChIP) assay was previously described (Cai *et al.*, 2014). The nucleus was isolated from 10-day-old wild type and *35S:HA-ERF109* seedlings and the chromatin was sonicated to fragments with various sizes (250 bp-1 kb). The DNA fragments bonded by HA-ERF109 were then enriched by anti-HA antibodies (1:100 for ChIP assay, HA-Tag, Mouse mAb, M20003, Abmart, Shanghai, China) from whole DNA fragment pool. After incubation with Protein A agarose beads (Millipore, USA), enriched DNA fragments used as the templates for PCR were eluted out and purified with phenol/chloroform (1:1, v/v). The enriched DNA fragments can be detected by PCR and quantified by quantitative-PCR with specific primers which were showed in Supplemental Table S1.

### ACCESSION NUMBER

Sequence data from this article can be found in the Arabidopsis Genome Initiative or GenBank/EMBL databases under the following accession numbers: *ERF109*, *At4g34410*; *PID*, *At2g34650*; *PIN1*, *AT1G73590*; *PIN2*, *At5g57090*; *PIN3*, *AT1G70940*; *PIN4*, *At2g01420*; *PIN7*, *AT1G23080*; *AUX1*, *AT2G38120*; *EDT1/HDG11*, *At1g73360*; *UBQ5*, *At3g62250*.

## SUPPORTING INFORMATION

Supplemental Figure S1. Expression level of auxin transport-related genes in *35S:ERF109* and wild type.

Supplemental Figure S2. Verification of *ERF109* expression level in the lines for genetic assay (*35S:ERF109 pid*, *35S:ERF109 pin2*, *35S:ERF109 pin4*).

Supplemental Table S1. Primers used in this study.

## ACKNOWLEDGEMENTS

This work was supported by grants from the National Nature Science Foundation of China (31572183) and the Ministry of Science and Technology of China (2016YFD0100701, 2018ZX08009-11B, 2016ZX08005-004-003, 2016ZX08001003). We thank Dr. Chuanyou Li, Institute of Genetics and Developmental Biology, CAS for providing PIN-GFP marker lines

## AUTHORS CONTRIBUTION

LR, XTC designed the research, performed the research, data analysis and interpretation.

PXZ, PX performed the research, data analysis and interpretation.

LR wrote the manuscript and XTC revised the manuscript.

CBX designed the research, performed interpretation, edited the manuscript, and supervised the project

